# Updated chromosome-level genome assembly of *Sporisorium scitamineum* with improved accuracy and completeness

**DOI:** 10.1101/2025.05.23.649816

**Authors:** Lucas M. Taniguti, Pedro F. Vilanova, João P. Kitajima, Claudia B. Monteiro-Vitorello

## Abstract

In 2015, we published the complete genome assembly of *Sporisorium scitamineum*, the fungal pathogen responsible for sugarcane smut disease, generated from PacBio long-read sequencing and polished with Illumina short reads. Since then, the tools for genome assembly have improved considerably, including advances in sequencing technologies and bioinformatics workflows, motivating us to revisit our original assembly. Here, we present an updated genome version, using newly generated high-quality Illumina short-read data and DeepVariant-based polishing, resulting in corrections at approximately 5,500 genomic positions. BUSCO benchmarking in protein mode demonstrated increased genome completeness from the original 94.2% to 99.3%, indicating substantial improvements in both accuracy and gene annotation reliability. Additionally, comparison with publicly available assemblies reveals that most sequence polymorphisms occur in isolates belonging to the Asian lineage, consistent with earlier population-level studies. This enhanced genome assembly provides a significantly improved reference resource, enabling more robust genetic, evolutionary, and functional studies of this economically important sugarcane pathogen.

## INTRODUCTION

Genome assembly involves organizing sequencing reads into a contiguous representation of an organism’s chromosomes by aligning and merging data from various sequencing platforms (Basantani et al. 2017; Baker 2012). Currently, sequencing technologies differ notably in read length, accuracy, and cost. Short-read platforms, such as Illumina and MGI, provide accurate yet shorter reads (Kim et al. 2021), whereas long-read platforms like PacBio and Nanopore generate reads spanning thousands of base pairs but with higher raw error rates (Jain et al. 2018; Karst et al. 2021). Hybrid assembly approaches leverage the strengths of different sequencing technologies, mitigating their individual limitations to enhance assembly accuracy (Chen et al. 2020).

*Sporisorium scitamineum* causes sugarcane smut disease, significantly reducing crop yield worldwide. Plants infected by this fungus typically show severe growth impairment, with reported yield losses ranging from 12% to as high as 75% (Rajput et al. 2024). Understanding the genome of this pathogen may reveal essential information on virulence, effector candidates, and host adaptation, guiding breeding for resistance and improving disease control strategies. Our initial genome assembly of *Sporisorium scitamineum*, published in 2015 (Taniguti et al. 2015), utilized PacBio long-read sequencing and Illumina short-read polishing. However, despite this hybrid approach, residual errors remained, potentially limiting downstream genetic and functional analyses.

Recent advancements in sequencing technologies and computational methods now allow for improved accuracy and completeness of genome assemblies. Short-read sequencing data from newer Illumina platforms, combined with variant-calling methods such as DeepVariant (Poplin et al. 2018), can identify and correct errors in reference assemblies, leading to a refined consensus sequence. Additionally, more comprehensive gene prediction pipelines like FunGAP (Min et al. 2017) integrate multiple gene prediction tools along with RNA sequencing (RNA-seq) data, improving on our previous approach that relied solely on Augustus (Stanke et al. 2006).

Recognizing these technological advances, we revisited our original *S. scitamineum* genome assembly. Here, we report an updated genome assembly obtained by polishing the original genome with newly generated Illumina sequencing data, variant calling, and consensus refinement techniques. We also improved gene prediction by utilizing the more complete FunGAP pipeline, incorporating existing RNA-seq data. Comparison of our updated assembly with other available *S. scitamineum*’s genomes demonstrated that the Chinese assembly (Que et al. 2014) had most of the variation, while South African assemblies shared a higher similarity. The resulting genome resource represents an advancement, facilitating more accurate genetic, evolutionary, and functional studies of this pathogen.

## METHODS

### DNA Extraction, Library Preparation, and Sequencing

Fungal cells from MAT-1 haploid yeast-like cells derived from SSC39 teliospores were grown in a liquid medium YM (3 g L^−1^ yeast extract, 3 g L^−1^ malt extract, 5 g L^−1^ peptone, 10 g L^−1^ glucose) at 28 °C with shaking at 200 rpm. Genomic DNA was extracted with a modified Doyle and Doyle protocol (Doyle and Doyle 1987) to meet sequencing quality standards.

The library preparation was then performed using the DNA Prep Kit (Illumina, San Diego, CA, USA). The resulting library was sequenced on a NextSeq 2000 System in 2 × 300 bp paired-end runs at the Functional Genomics Center, ESALQ, University of São Paulo, Piracicaba, Brazil.

### RNA data

RNA data were obtained from a previous study (Taniguti et al. 2015), in which haploid yeast-like cells of opposite mating types were grown separately in a liquid medium for 15 h at 28 °C under constant shaking. The cells were then mixed before RNA extraction, flash-frozen in liquid nitrogen, and stored at −80 °C until use. Paired-end sequencing was performed on the Illumina HiScanSQ platform.

### Polishing

The original reference genome for *S. scitamineum* (NCBI accession GCA_001010845.1) (Taniguti et al. 2015) was retrieved from GenBank. Illumina paired-end reads generated in the present study were aligned to this assembly with `bwa-mem` (Li 2013). Variants were then called using `Deep-Variant` (v1.6.0, model WGS) (Poplin et al. 2018), and the resulting VCF file was applied to the reference with `bcftools consensus` (Danecek et al. 2021) to create a polished FASTA sequence (deposited under accession GCA_001010845.3).

#### Assembly evaluation

Completeness and basic contiguity statistics were obtained by running `busco` v5.7.0 (Manni et al. 2021) in genome mode (`-m genome`) with the *basidiomycota_odb10* lineage dataset and `AUGUSTUS` as the internal gene predictor. The same BUSCO run and a `quast` v5.2 analysis (default settings) were applied to five additional publicly available *S. scitamineum* assemblies (GCA001010845.1, GCA928991175.1, GCA000772675.1, GCA900002365.1, and GCA023212615.1) to ensure that all genomes were evaluated under identical conditions. For each assembly, the following metrics were extracted from tool outputs: number of contigs, total assembly length, proportion of ambiguous bases (% gaps), GC content, and *N*_90_.

### Whole-genome alignment and variant calling

For each sample, the query assembly was aligned to the resulting reference genome using MUMmer v4.0.0-rc1 (`nucmer`, default parameters), and filtered to retain only one-to-one matches with `delta-filter −1`; the resulting `*.delta` file was converted to VCF with `delta2vcf`, after which two in-place edits with `sed` command to correct the genotype field and prune superfluous FORMAT sub-fields.

### Gene Prediction

Gene models were generated with the FunGAP pipeline, which combines multiple *ab initio* predictors (`AUGUSTUS`, `SNAP`, `GeneMark-ES`) and integrates RNA-seq evidence to refine exonintron boundaries.

#### Proteome evaluation

The translated protein set produced by FunGAP was evaluated with `busco` v5.7.0 in protein mode (`-m proteins`) against the same *basidiomycota_odb10* dataset to verify the completeness of the conserved fungal orthologues at the annotation level.

## RESULTS AND DISCUSSION

### Sequencing and genome polishing

Illumina paired–end sequencing generated 1.59 Gbp of data (2 × 300 bp), corresponding to roughly 80x coverage of the 20 Mb genome. Read quality profiles for R1 and R2 were nearly identical, with an average GC content of 53%, a sharp length distribution (290-300 bp) and minimal adaptor or low–quality sequence removal.

Alignment of these reads to the 2015 reference followed by `DeepVariant` calling introduced 5,500 single-nucleotide and indel corrections. The resulting polished assembly, GCA_001010845.3, spanned 20.07 Mb across 27 contigs and was completely gap-free.

### Assembly quality compared with public references

The new assembly presented an *N*_90_ of 573 kb, the highest among the six assemblies we benchmarked (Table1). Genome-mode BUSCO v5.7.0 recovered 98.8 % complete orthologues, surpassing the original 2015 build (96.5 %) and marginally outscoring the next–best public assembly (98.7 %). The absence of gaps ensures uninterrupted representation of repeat–rich regions, including the mating–type loci that underpin pathogenicity in smut fungi.

**TABLE 1.**
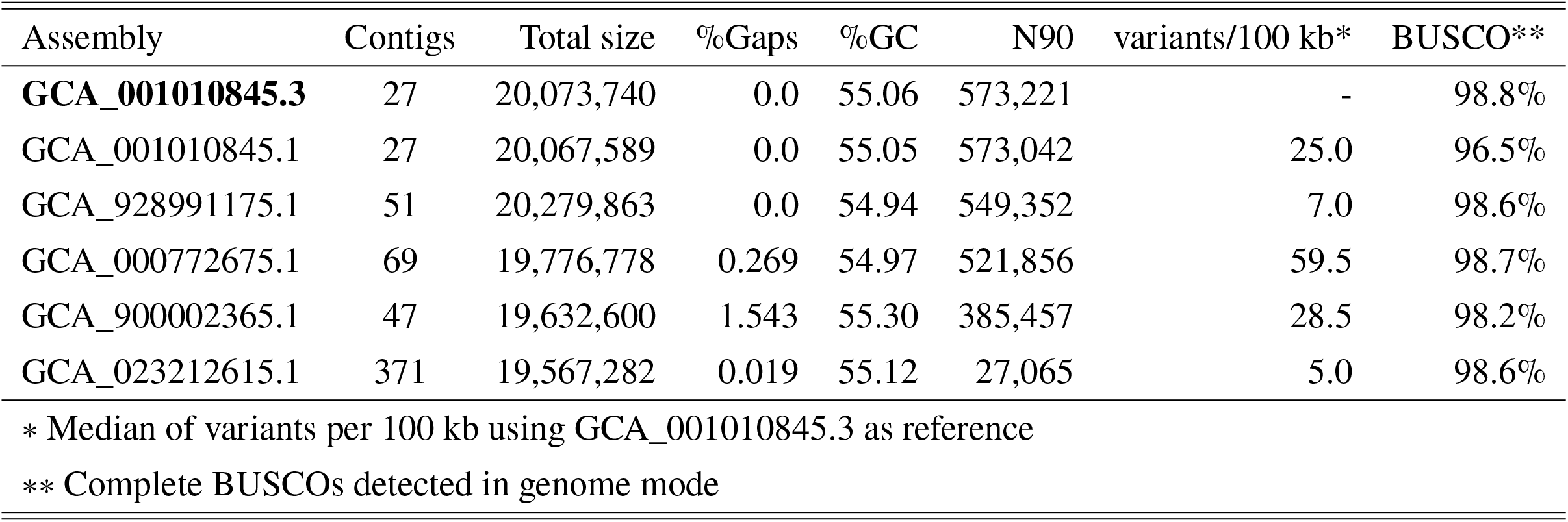
Comparative assembly statistics for the new polished genome versus five previously published S. scitamineum references.

#### Small-variant density

Using our updated reference genome (GCA_001010845.3) as the baseline, we detected 25 single-nucleotide and small-indel variants per 100 kb when the original 2015 assembly (GCA_001010845.1) was realigned (Table 1). Even assemblies from other geographic origins show fewer differences relative to the new reference than did our previous draft, underscoring the extent of the corrections introduced during polishing. Among publicly available genomes, the Chinese isolate GCA_000772675.1 was the most divergent, with median of 59.5 variants per 100 kb. This pattern is consistent with population-level studies indicating that American and African isolates derive from a single Asian lineage, reflecting a founder effect (Raboin et al. 2007). Most other assemblies also represent MAT-2 strains, whereas our reference is MAT-1, which may contribute to the observed differences. Finally, variation attributable to reference choice, alignment parameters, repeat masking, sequencing depth and quality, and platform-specific biases should be considered when interpreting absolute variant counts (Zverinova and Guryev 2022).

### Gene prediction and proteome completeness

FunGAP predicted 6,907 protein–coding genes with a mean CDS length of 1,761bp (median 1,425bp). Approximately 31 % of genes contained at least one intron, and the average protein length was 587 amino acids. Proteome–mode BUSCO identified 99.3 % complete conserved orthologues, confirming that nearly the entire fungal gene complement is captured and correctly modelled. This represents a gain of five percentage points over the 2015 annotation.

### Reproducibility

Every step of the analysis pipeline—including read mapping, variant calling, consensus generation and benchmarking—is encapsulated in the publicly archived workflow `AssemblyPolish.wdl`. Container images and parameter files are version–controlled, enabling any researcher to reproduce the results on local hardware or in the cloud.

## CONCLUSIONS

Taken together, these gains in sequence accuracy, contiguity, and gene-space completeness position GCA001010845.3 as the most accurate and comprehensive reference genome available for *S. scitamineum* to date. This resource will facilitate fine-scale variant discovery, allele-specific expression analyses, and high-resolution comparative genomics across smut fungi. In addition, this genome-wide variation assessment confirmed previous reports of lower genetic diversity among American and African isolates compared to Asian lineages. These insights shed light on population diversification at genome-level and lay the groundwork for future studies focused on understanding pathogen evolution.

## Supporting information

Gene annotations in GFF3 format

FASTA with aminoacid sequences

BUSCO result for protein mode

BUSCO result for genome mode

New version of genome in FASTA format

WDL release with workflow used for polishing

## DATA AVAILABILITY

The polished *Sporisorium scitamineum* genome generated in this study is publicly available in the NCBI Nucleotide database under chromosome-level accessions CP010913.2-CP010939.2 (assembly accession `GCA_001010845.3`).

To maximize reproducibility and make the new gene predictions accessible, we provide the following items as supplementary material.

**Polished genome sequence**: the exact gap-free FASTA file corresponding to the public NCBI assembly.

**Gene predictions**: a comprehensive `.gff` file containing coordinates and features for all 6 907 predicted genes, plus a companion `.faa` file with the translated amino acid sequences of every coding region, providing a ready resource for downstream studies.

**Polishing workflow**: the complete pipeline used to generate the final assembly.

**BUSCO completeness reports**: original outputs for both *genome* and *protein* modes (v5.7.0, *basidiomycota_odb10* dataset).

The `AssemblyPolish.wdl` workflow is also mirrored on GitHub. All other supplementary files accompany this manuscript.

## FUNDING

The authors acknowledge the support of the Brazilian institution FAPESP: grant numbers 2022/03962-7 and CNPq 405314/2021-3 and 305961/2017-7 (C.B.M.-V.); fellowships to P.F.V. FAPESP (2023/13474-2).

Mendelics Análise Genômica also provided support in the form of a salary for authors L.M.T. and J.P.K.

These sponsors had no role in study design, data collection and analysis, decision to publish or manuscript preparation.

## ACKNOWLEDGEMENTS

The authors thank Elaine Vidotto Batista for technical support and Marcella Ferreira (Microbe Lab, ESALQ/USP) for participating in DNA extraction.

