## Supplementary material for "Updated chromosome-level genome assembly of *Sporisorium scitamineum* with improved accuracy and completeness": BUSCO result for protein mode

# BUSCO version is: 5.7.0 
# The lineage dataset is: basidiomycota_odb10 (Creation date: 2024-01-08, number of genomes: 133, number of BUSCOs: 1764)
# Summarized benchmarking in BUSCO notation for file /mnt/miniwdl_task_container/work/_miniwdl_inputs/0/SSC39-MAT1.polished.renamed.faa
# BUSCO was run in mode: proteins

	***** Results: *****

	C:99.3%[S:99.3%,D:0.0%],F:0.3%,M:0.4%,n:1764	   
	1752	Complete BUSCOs (C)			   
	1752	Complete and single-copy BUSCOs (S)	   
	0	Complete and duplicated BUSCOs (D)	   
	6	Fragmented BUSCOs (F)			   
	6	Missing BUSCOs (M)			   
	1764	Total BUSCO groups searched		   

Dependencies and versions:
	hmmsearch: 3.1
	python: sys.version_info(major=3, minor=7, micro=12, releaselevel='final', serial=0)
	busco: 5.7.0
