## Supplementary material for "Updated chromosome-level genome assembly of *Sporisorium scitamineum* with improved accuracy and completeness": WDL release with workflow used for polishing: AssemblyPolish.wdl

```
# This workflow performs assembly polishing using Illumina sequencing data.
#
# It includes the following steps:
# 1. Genotyping with Illumina data to identify variants.
# 2. Generating a consensus sequence from the identified variants.
# 3. Re-aligning Illumina data to the consensus sequence to obtain updated alignments and variants.
# 4. Assessing the polished assembly using BUSCO.

version development-1.1

import "structs.wdl"
import "bcftools_consensus.wdl"

import "IlluminaGenotyping.wdl" as illumina_genotyping
import "AssemblyAssessment.wdl" as assembly_assessment

workflow AssemblyPolish {
  input {
    PairedEndExperiment illumina_dna_experiments  # Suport only one experiment
    File reference_genome
    String output_name
    File busco_dataset
    String lineage_name
    Int threads = 8
    String program = "deepvariant"
    Boolean stub = false
  }


  call illumina_genotyping.IlluminaGenotyping {
    input:
      illumina_dna_experiments = [illumina_dna_experiments],
      reference_genome = reference_genome,
      threads=threads,
      program=program,
      stub = stub
  }


  call bcftools_consensus.Consensus {
    input:
      basename = output_name,
      fasta = reference_genome,
      variants = IlluminaGenotyping.variants[0],
      stub = stub
  }

  # Need only new alignment. Conveniente use Genotyping pipeline because of reports
  call illumina_genotyping.IlluminaGenotyping  as final {
    input:
      illumina_dna_experiments = [illumina_dna_experiments],
      reference_genome = Consensus.consensus,
      threads=threads,
      stub = stub
  }

  # Assessing
  call assembly_assessment.AssemblyAssesment {
    input:
      genome = Consensus.consensus,
      busco_dataset = busco_dataset,
      lineage_name = lineage_name,
      stub = stub
  }

  output {
    File consensus = Consensus.consensus
    File report = final.multiqc_report
    File report_raw = final.multiqc_raw_report
    File alignment = final.alignments[0]
    File alignment_idx = final.alignment_indices[0]
    File variants = final.variants[0]
    File variants_idx = final.variant_indices[0]
    Array[File] busco_genome = AssemblyAssesment.genome_outputs
    Array[File]? busco_proteins = AssemblyAssesment.protein_outputs
    Array[File] versions = flatten([final.software_versions, [Consensus.version, AssemblyAssesment.software_version]])
  }
}
```
